# Asian *Mus musculus*: Subspecies Divergence, Genetic Diversity, and Historical Biogeography

**DOI:** 10.1101/2024.03.19.585722

**Authors:** Yaser Amir Afzali

## Abstract

The house mouse, *Mus musculus*, is a model organism that has greatly contributed to evolutionary research. Despite its significance, there remain gaps in our understanding of its phylogeography and population genetic structure in Asian regions. This comprehensive study, aims to elucidate the evolutionary history, genetic diversity, and distribution patterns of *M. musculus*. A diverse dataset of 281 *M. musculus* samples from across Asia was collected, covering 101 localities in 19 countries. Phylogenetic analysis using Cytochrome *b* (*Cytb*) and D-Loop (*DL*) region, unveiled well-supported lineages. These lineages correspond to four major subspecies: *Mus musculus musculus*, *M. m. domesticus*, *M. m. castaneus*, and *M. m. bactrianus*. The analysis suggests a monophyletic origin of these subspecies approximately 0.59 million years ago (Mya), followed by two main lineages: one consisting of *M. m. domesticus* (∼0.59 Mya) and the other comprising *M. m. castaneus*, *M. m. bactrianus*, and *M. m. musculus* (∼0.56 Mya). Genetic diversity varied among subspecies, with *M. m. domesticus* exhibiting the highest diversity due to its extensive global distribution and *M. m. bactrianus* exhibiting the lowest diversity due to restriction in the southwest Asia. Pairwise genetic distances and *FST* values confirmed significant genetic differentiation among the subspecies, underlining their historical isolation. Additionally, a Mantel test indicated that geographical distance played a pivotal role in shaping genetic differentiation. Demographic analysis revealed evidence of population expansions in *M. m. domesticus*, *M. m. musculus*, and *M. m. castaneus*, while *M. m. bactrianus* showed characteristics of neutral selection and genetic drift. Species distribution modeling, assessing both Current Conditions and the Last Glacial Maximum, indicated habitat shifts and losses during glacial periods, particularly in the eastern and northern regions of Asia. However, each subspecies displayed unique responses to climatic changes, reflecting their distinct ecological adaptations. Historical biogeography analysis pointed to complex origins and a network of dispersal and vicariance events that led to the contemporary distribution of subspecies. Deserts and Xeric Shrublands emerged as critical areas for diversification and speciation. This study contributes to our understanding of the phylogeography and population genetics of *M. musculus* in Asia, highlighting the significance of geographical factors and climatic fluctuations in shaping its evolutionary history and genetic diversity.

## INTRODUCTION

The house mouse, scientifically known as *Mus musculus* Linnaeus, 1758, has long served as an invaluable model for evolutionary research, with its genome among the first to be fully sequenced (Church et al., 2009). Previous molecular investigations have identified five major subspecies: *Mus musculus musculus* Linnaeus, 1758 in Eastern Europe, Central and North East Asia (Eastern European house mouse); *Mus musculus domesticus* Schwarz and Schwarz, 1943 in Western Europe, North America, South America, Africa and Oceania (Western European house mouse); *Mus musculus castaneus* Waterhouse, 1843 in southern and southeastern Asia (Southeastern Asian house mouse); *Mus musculus bactrianus* Blyth, 1846 in southwestern and Central Asia (Southwestern Asian house mouse) and *Mus musculus gentilulus* Thomas, 1919 in the Arabian Peninsula and Madagascar (pygmy house mouse) (Mitchell-Jones et al., 1999; Musser & Carleton, 2005; Darvish et al., 2012; Amir Afzali et al., 2017; Amir Afzali, 2024). These subspecies have been supported as distinct evolutionary lineages through various studies and methodologies (Sakai et al., 2005; Frazer et al., 2007; Suzuki et al., 2013; Li et al., 2021). However, while *Mus musculus* is a well-studied species due to its prominent role as a laboratory model, our understanding of its phylogeography and population genetic structure remains incomplete (Guénet & Bonhomme, 2003; Hardouin et al., 2015; Maltsev et al., 2016; Amir Afzali & López-Antoñanzas, 2024). Over the past two decades, the field of phylogeography, which explores the geographical distribution of genealogical branches within closely related species, has gained prominence (Avise, 2000; Maltsev et al., 2016; Amir Afzali et al., 2018; Amir Afzali et al., 2024). This approach allows us to fill gaps in the history of *M. musculus* distribution, examine its evolutionary history, and understand its population genetic structure in the context of geographical barriers, historical geological and climatic events, and speciation patterns. In this study, we aim to achieve several objectives: I) explore *M. musculus*’s evolutionary history using mitochondrial markers; II) establish a time-calibrated phylogenetic hypothesis; III) assess the impact of geographical barriers, historical geological, and climatic events on the population genetic structure; IV) evaluate the effects of Pleistocene climatic fluctuations; and V) contribute to the broader understanding of *M. musculus*.

## MATERIAL AND METHODS

### Sample Collection

In this study, a comprehensive dataset of 281 *M. musculus* samples were collected, encompassing representatives from four distinct subspecies of house mice in Asia. Sampling was expansive, spanning 101 localities distributed across 19 countries (SD1). This geographic range covered over 23 million km^2^, ensuring that a broad representation of the species’ distribution within the region (Fig. 1). The selection of sampling localities aimed to achieve maximal coverage of the species’ geographic range and included the most taxonomically informative sampling within the study area. This approach allowed us to obtain a diverse and representative set of samples for our analysis.

**Fig. 1.**
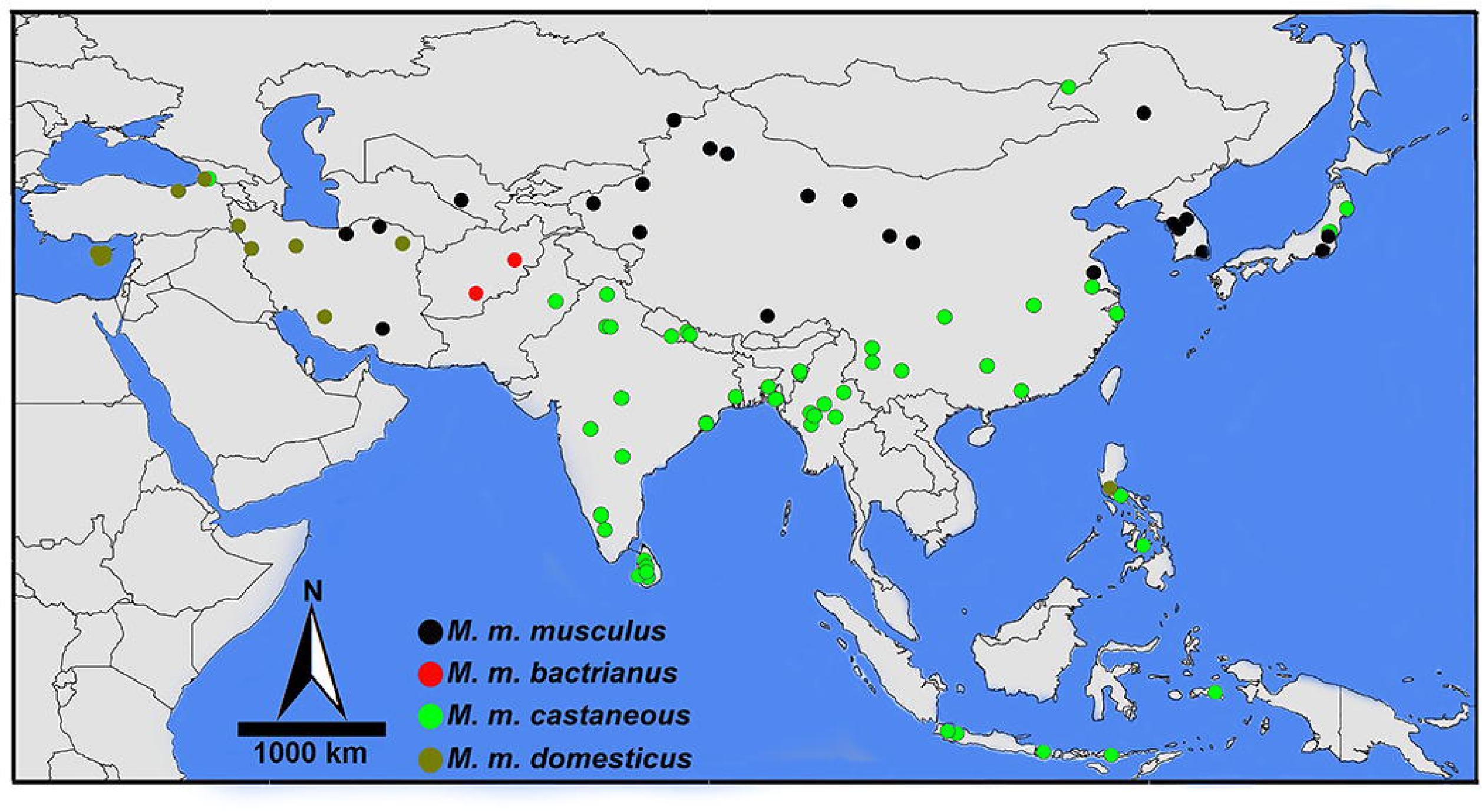
Sampling localities of *Mus musculus* in Asia.

### DNA Extraction and PCR Amplification

Total genomic DNA was extracted from muscle tissue samples using the Sambio DNA reagent kit, following the manufacturer’s instructions. Two mitochondrial regions targeted for amplification: the Cytochrome *b* gene (*Cytb*) and the D-Loop (*DL*) region. For the *Cytb* gene, we used the L7 and H6 primers, following the protocol described by Montgelard et al. (2002). The amplification of the D-Loop (*DL*) was conducted using the CR1 and CR2 primers. These primers have been previously utilized in studies by Yasuda et al. (2005) and Suzuki et al. (2013). PCR amplification for both the *Cytb* and *DL* followed previously established protocols. The reaction conditions included an initial denaturation step, typically at 95°C for 3 minutes, followed by a specific number of amplification cycles, each consisting of denaturation at 94°C for 45 seconds, annealing at 55°C for 45 seconds, and extension at 72°C for 60 seconds. A final extension step was performed at 72°C for 7 minutes. The obtained sequences were subjected to manual editing for quality control. We used ChromasPro version 2.6.6 (http://technelysium.com.au) and BioEdit (version 7.0.5.3; Hall, 1999) to ensure high-quality sequences. Alignment of sequences was carried out using the MUSCLE algorithm with default parameters within MEGA-X (Kumar et al., 2018). All sequences generated during this study have been deposited in the GenBank database. To expand the breadth and depth of our dataset, the newly generated sequences were merged with available data from GenBank (see SD2, SD3 and SD4). This comprehensive approach ensured that our analyses encompassed a diverse and informative set of sequences, spanning a wide geographic range within the study area.

### Phylogenetic Analyses

We conducted phylogenetic analyses using two distinct approaches: Maximum Likelihood (ML) with RAxML 8.2.12 (Stamatakis, 2014) and Bayesian Inference (BI) with BEAST 2 version 2.7.3 (Bouckaert et al., 2019). The appropriate models of nucleotide substitution were selected with jModelTest 2.1.10 (Darriba et al., 2012) using the Akaike Information Criterion (AIC). For the Bayesian Inference analysis, four Markov chain Monte Carlo (MCMC) simulations were run for 40 million generations, with samples collected at every 10,000th tree. Convergence of the MCMC runs was assessed using Tracer v1.7.0. All parameters’ effective sample sizes were verified to exceed 200, ensuring the reliability of the results. The maximum clade credibility tree (MCC) was derived and annotated using TreeAnnotator 2.7.3, following the removal of the initial 25% of trees as burn-in. To estimate the maximum-likelihood tree, we conducted analyses within RAxML 8.2.12 and performed 1,000 bootstrap replications to assess the robustness of the tree. The resulting trees were visualized using FigTree v1.4.4 software (Rambaut, 2018).

### Divergence Time Analysis

To generate a time-calibrated phylogenetic topology, we used the uncorrelated relaxed log-normal clock model within BEAST 2 version 2.7.3 (Bouckaert et al., 2019). In this analysis, we used *Mus macedonicus* Petrov & Ruzic, 1983, and *Mus spicilegus* Petényi, 1882, as outgroup taxa to root the tree. A calibration point was applied based on a previously established value of 0.91 million years ago (Mya) for the most recent common ancestor (MRCA) of *M. macedonicus* and *M. spicilegus*. This calibration point was sourced from Suzuki et al. (2004, 2013). MCMC simulations were executed with a total of 40 million generations, with samples collected every 10,000 iterations. We assessed the convergence of the MCMC run, ensuring that all parameters had an effective sample size exceeding 200, to ensure the robustness and reliability of the results. This assessment was performed using Tracer v1.7.0. After the MCMC run, the initial 25% of trees were discarded as burn-in. Subsequently, a maximum clade credibility tree was generated. The resulting trees were visualized and interpreted using FigTree v1.4.4 (Rambaut, 2016).

### Population Structure

To assess population structure, we calculated standard genetic diversity metrics, which included nucleotide diversity (π), haplotype diversity (Hd), the number of haplotypes (h), and the number of segregating sites (s) using DnaSP 6.12.03 (Rozas et al., 2017). To visualize the relationships among haplotypes, we constructed haplotype networks using statistical parsimony network analysis, specifically the TCS Network, implemented in PopART (Clement et al., 2002). We assessed genetic differentiation and structure among lineages using pairwise *FST* values in Arlequin v3.5.2.2 (Excoffier & Lischer, 2010). This analysis provided insights into the genetic divergence between populations. Mean genetic divergences among lineages were calculated using the p-distance model with 1,000 bootstrap replicates in MEGA-X (Kumar et al., 2018). This approach allowed us to quantify the genetic differences between lineages and populations. To explore the relationship between geographical distance and genetic distance among *M. musculus* populations, we conducted a Mantel test using Arlequin v. 3.5.2.2. This analysis helped us evaluate the extent to which geographical factors influence genetic differentiation.

### Historical Demography

We used Arlequin v3.5.2.2 to conduct neutrality tests and mismatch distribution (MMD) analysis. These analyses aimed to identify any deviations from the null hypothesis of neutral evolution and to assess recent population expansion. Each analysis involved 10,000 permutations to ensure robust results. To evaluate deviations from the null hypothesis of a neutral model of evolution, we calculated neutrality indices, specifically Tajima’s D and Fu’s F statistics for each lineage. These indices were calculated following the approach outlined by Excoffier and Lischer (2010). The onset of population expansion in all groups was estimated using the formula: t = τ/2uk, where t represents the time since expansion, τ is the expansion parameter tau, u is the evolutionary rate per generation, and k is the sequence length. This approach, as described by Rogers and Harpending (1992) and Rogers (1995), allowed us to estimate the time since population expansion. The expansion parameter tau (τ) was estimated for the *Cytb* sequences using Arlequin v3.5.2.2. Three classes of τ values were identified for the *Cytb* sequence analysis: small (τ ∼ 2–4), intermediate (τ ∼ 5–6), and large (τ ∼ 7–9). Based on these τ values, we applied three different evolutionary rates: 0.11, 0.047, and 0.028 substitutions per site per million years (substitutions/site/myr) for calculating the time since expansion. These rates were adapted from Honda et al. (2019) and Maung Maung Theint et al. (2021) to estimate the expansion parameters effectively.

### Species Distribution Modelling

We utilized Maxent 3.4.4 (Phillips et al., 2006) to estimate the probability of species occurrence. To perform this analysis, we used presence data in conjunction with environmental variables, following the methodology outlined by Elith et al. (2006), Hernandez et al. (2006), and Rhoden et al. (2017). Nineteen bioclimatic variables were obtained from the WorldClim series (Hijmans et al., 2005; http://www.worldclim.org) to characterize the environmental conditions for our species distribution models. We ran the models for two temporal conditions: the Last Glacial Maximum (LGM) and Current Conditions (CC). The spatial resolution was set at 2.5 arc-minutes (approximately 5 km^2^) for LGM and 30 arc-seconds (approximately 1 km^2^) for CC. In model training, %75 of the presence records was used, while %25 was randomly selected for testing. The procedure was repeated 15 times, with 5,000 iterations performed. The accuracy of the models was evaluated using receiver operating characteristic (ROC) analysis. The Area Under the Curve (AUC) derived from the ROC plot, ranging between 0 and 1, served as the key metric. A model with an AUC value greater than 0.75 was considered robust and acceptable, while AUC values below 0.5 indicated a random prediction (Elith et al., 2006). To determine the importance of each climatic variable in explaining the species distribution, we conducted a jackknife procedure, following the approach outlined by Sillero and Carretero. (2012). This procedure allowed us to identify the key environmental variables driving the observed distribution patterns.

### Historical Biogeography

We conducted ancestral biogeography reconstruction using the Dispersal-Extinction-Cladogenesis (DEC) model within the BioGeoBears package (Ree et al., 2005; Matzke 2013; Ree & Sanmartín, 2018). From the time-calibrated tree obtained through Bayesian Inference analyses in BEAST 2 version 2.7.3 (Bouckaert et al., 2019), we extracted the ingroup subtree. To estimate the posterior probabilities of ancestral areas at each node, we used a total of 3,604 trees (after discarding the initial 25% burn-in). This analysis allowed us to infer the likely distribution areas at various points in the evolutionary history of *M. musculus* in Asia. The distribution range of *M. musculus* in Asia was classified into eight major biogeographic areas based on the framework defined by Olson et al. (2001): A, Broadleaf and Mixed Forests; B, Grasslands, Savannas, and Shrublands; C, Deserts and Xeric Shrublands, D, Montane Grasslands and Shrublands, E, Moist Broadleaf Forests, F, Mediterranean Forests, Woodlands, and Scrub; G, Dry Broadleaf Forests; H, Coniferous Forests.

## RESULTS

### Phylogeographic Pattern

We sequenced 920 base pairs (bp) of the *Cytb* gene from 281 individuals and 875 bp of the *DL* from 51 individuals. Our phylogenetic analyses were based on a concatenated dataset comprising 1975 bp. Using the Akaike Information Criterion (AIC), we identified the best-fit models for the *Cytb* and *DL* datasets as GTR+I +G and HKY+I+G, respectively (Tavaré, 1986; Hasegawa et al., 1985). Both Maximum Likelihood (ML) and Bayesian Inference (BI) analyses yielded similar topologies. These topologies featured well-supported clades that corresponded with haplotype networks. Differences between the two analyses primarily lay in the level of clade support, which was generally higher in the BI analysis (SD5 and SD6). Our analyses indicated that all subspecies formed a monophyletic group. The MRCA of these subspecies is estimated to have originated approximately 0.59 million years ago (Mya). Subsequently, this MRCA diverged into two primary lineages. One lineage gave rise to the *M. m domesticus* (DOM), which originated around 0.59 Mya. The other lineage led to the *M. m. castaneus* (CAS), *M. m. bactrianus* (BAC) and *M. m. musculus* (MUS), with an estimated origin at about 0.56 Mya. Within this latter lineage, CAS, BAC, and MUS were identified as well-supported clades. CAS emerged approximately 0.37 Mya, BAC around 0.1 Mya, and MUS at about 0.29 Mya (Fig. 3). Importantly, all subspecies received strong support in both the BI and ML analyses, providing robust confirmation of their phylogenetic relationships and evolutionary history.

**Fig. 2.**
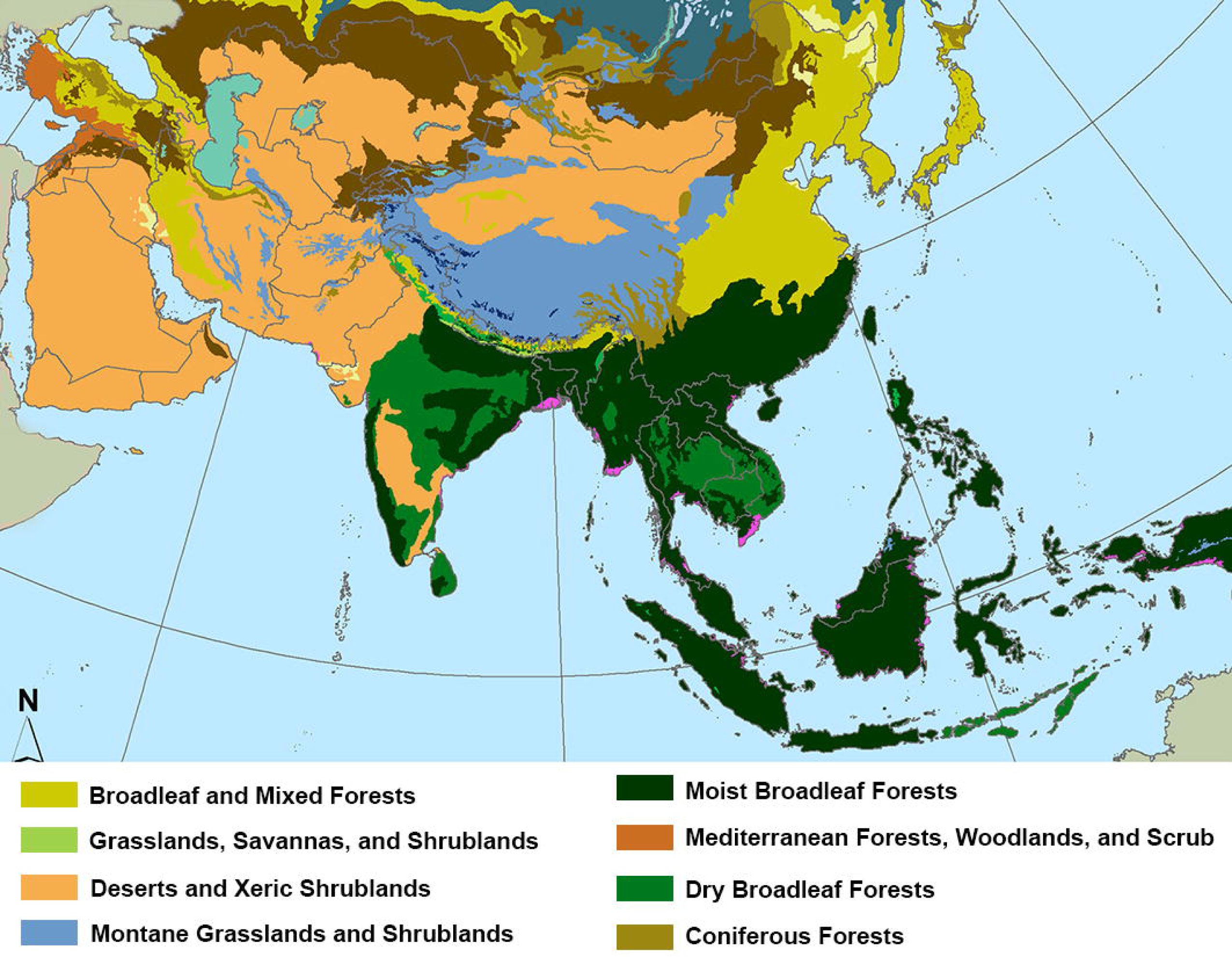
Terrestrial ecoregions in the Asia.

**Fig. 3.**
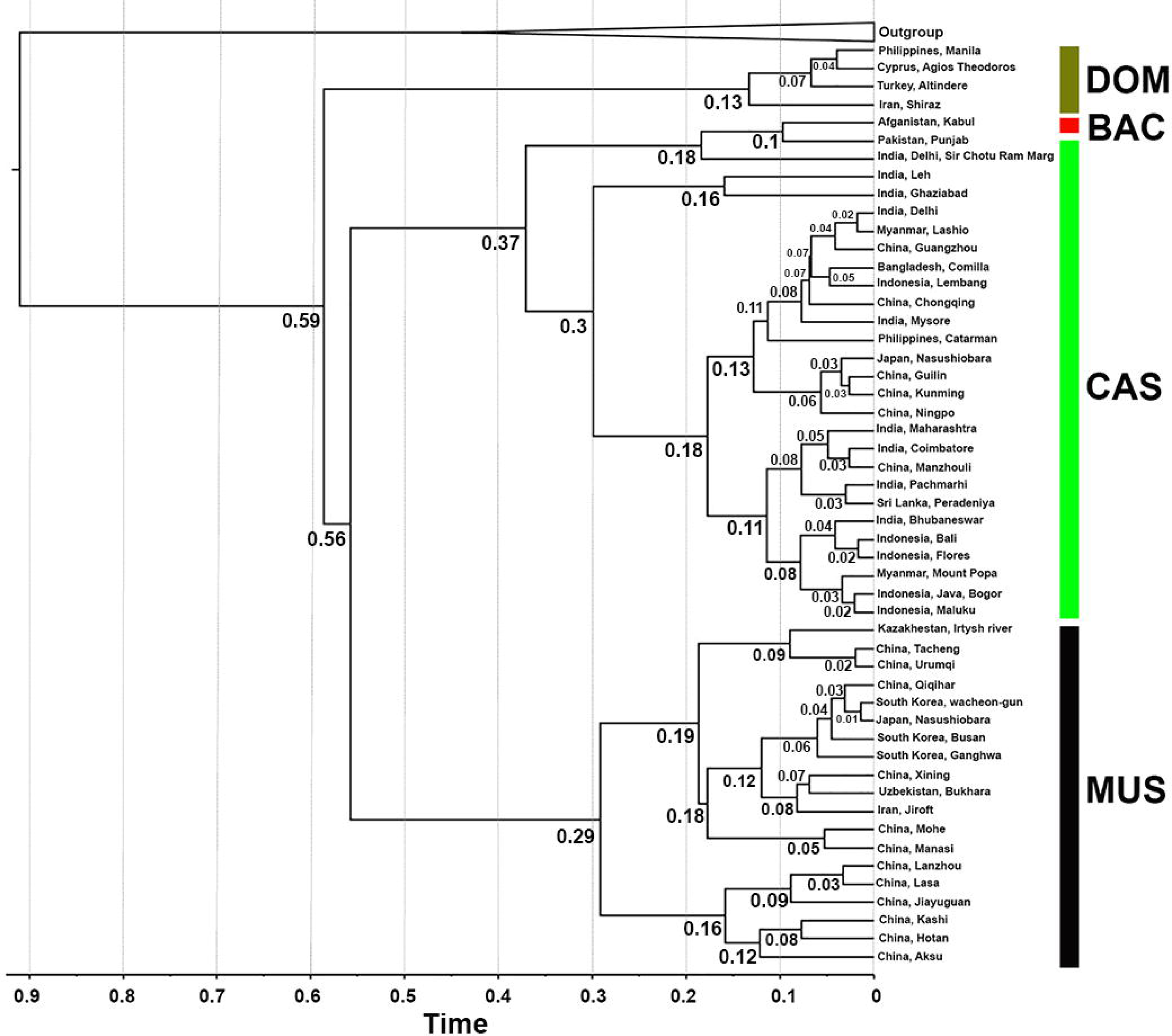
Bayesian Maximum Clade Credibility tree based on a concatenated dataset of *Cytb* and *DL* sequences. The numbers associated with each node indicate the estimated age of the respective node in million years ago (Mya).

### Genetic Diversity and Structure

The 920 bp *Cytb* fragment comprised 774 monomorphic sites, 146 polymorphic sites, 46 singleton sites, 100 parsimony informative sites, and a total of 151 mutations. These genetic variations defined 113 unique haplotypes within the 281 sampled specimens. Haplotype networks constructed for *Cytb* (Fig. 4) unveiled a distinct geographic pattern in the distribution of haplotypes. Notably, each subspecies exhibited its own exclusive haplogroups, with no sharing of haplotypes. Furthermore, within each subspecies, most populations were characterized by a single exclusive haplotype. The haplotype network revealed a clear pattern of genetic differentiation. CAS and BAC were positioned on one side, while MUS and DOM occupied the other side of the network. The highest genetic differentiation was observed between these two distinct sides of the haplotype network.

**Fig. 4.**
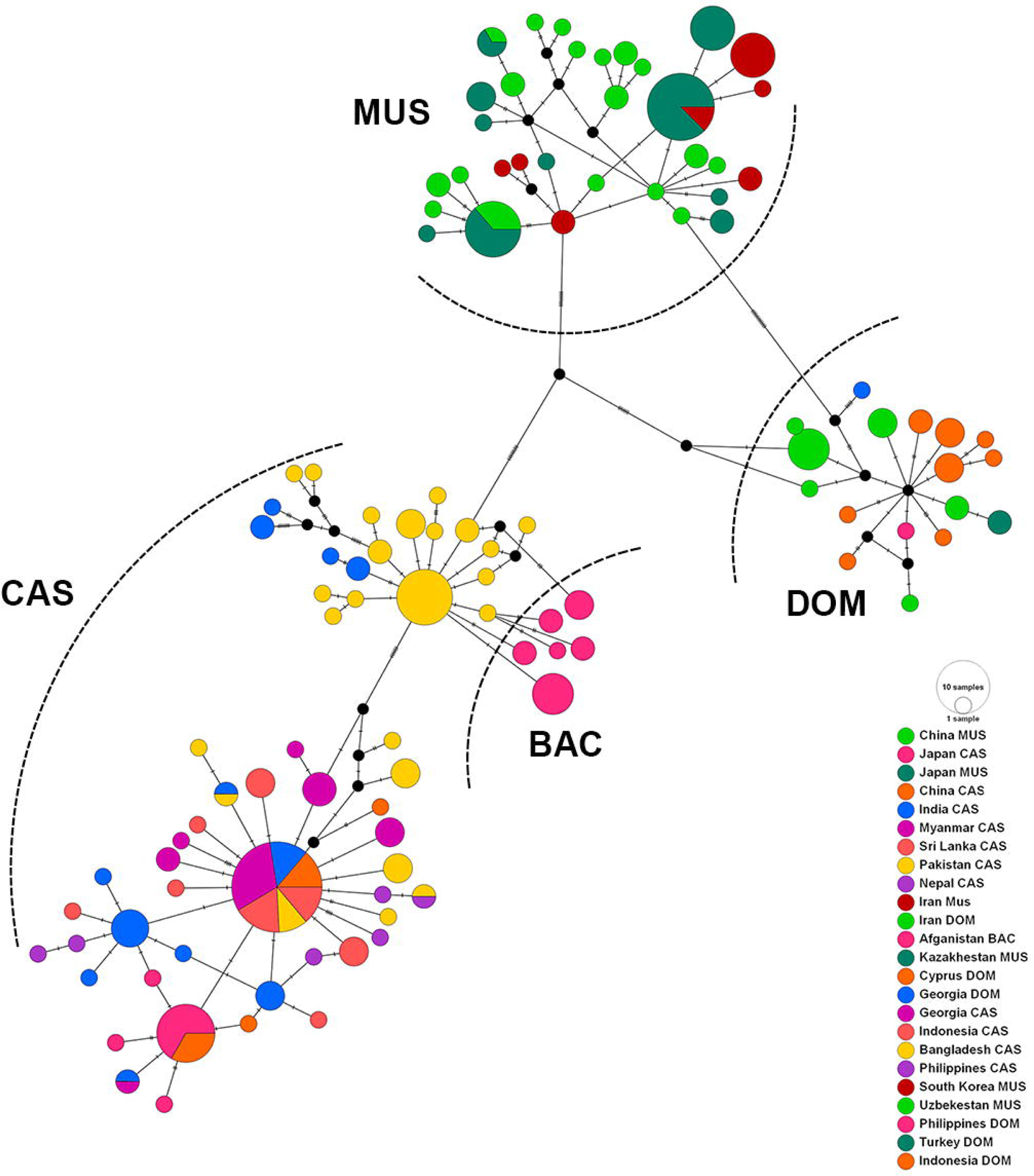
A 95% TCS network illustrating the relationships between the 113 *Cytb* haplotypes. The lines between circles in the network represent the number of mutational steps required to connect the haplotypes. This network provides a visual representation of the genetic relationships and divergence patterns among the haplotypes.

Considering the *Cytb* sequences, the overall mean genetic divergence across all subspecies was %2. Pairwise genetic divergence values are presented in Table 2, providing a detailed assessment of the genetic differentiation among *Mus musculus* subspecies.

**Table 1.** Genetic diversity indices of *Mus musculus*. Sample sizes (n), number of haplotypes (h), haplotype diversity (Hd), nucleotide diversity (π), and number of segregating sites (s).

|  | **n** | **h** | **Hd** | **$\pi$** | **s** |
| --- | --- | --- | --- | --- | --- |
| **MUS** | 90 | 29 | 0.91 | 0.00 | 35 |
| **DOM** | 37 | 16 | 0.89 | 0.00 | 31 |
| **CAS** | 138 | 51 | 0.88 | 0.00 | 71 |
| **BAC** | 16 | 6 | 0.82 | 0.00 | 10 |
| **ALL** | 281 | 113 | 0.97 | 0.01 | 146 |

**Table 2.** Mean genetic divergence based on *Cytb* between *Mus musculus* subspecies. Numbers are in percent.

|  | **DOM** | **MUS** | **CAS** | **BAC** |
| --- | --- | --- | --- | --- |
| **DOM** | * |  |  |  |
| **MUS** | 2 | * |  |  |
| **CAS** | 3 | 3 | * |  |
| **BAC** | 3 | 3 | 1 | * |

Pairwise *FST* comparisons based on *Cytb* sequences revealed significant genetic differentiation among all subspecies, as detailed in Table 3.

**Table 3.** Genetic distance *FST* between *Mus musculus* subspecies in Asia. Significant p values indicated with *.

|  | **DOM** | **MUS** | **CAS** | **BAC** |
| --- | --- | --- | --- | --- |
| **DOM** |  | * | * | * |
| **MUS** | 0.82 |  | * | * |
| **CAS** | 0.76 | 0.78 |  | * |
| **BAC** | 0.84 | 0.84 | 0.42 |  |

AMOVA tests conducted on the *Cytb* data revealed that genetic variation was distributed between subspecies (*FCT*) and among populations within subspecies (*FSC*). Notably, within each subspecies, the genetic variation was entirely distributed within populations (*FST*), as summarized in Table 4.

**Table 4.** AMOVA results of *Cytb* for *Mus musculus* in Asia. Significant p values indicated with *.

| Source of variation | d.f | Sum of squares | Variance components | Percentage of Variation | Fixation index |
| --- | --- | --- | --- | --- | --- |
| Among subspecies | 3 | 1674.34 | 9.05 | 74.82 | $F_{CT}=0.74^*$ |
| Among populations within subspecies | 19 | 286.88 | 1.17 | 9.73 | $F_{SC}=0.38^*$ |
| Within subspecies | 258 | 482.84 | 1.87 | 15.45 | $F_{ST}=0.84^*$ |
| Total | 280 | 2444.06 | 12.10 | 100 |  |

The Mantel test unveiled a positive and significant correlation between genetic and geographical distances, with a correlation coefficient (r) of 1.00 and a p-value of 0.00. This indicates that as geographical distances increase, genetic distances also significantly increase among subspecies.

### Historical Demography

Fu’s Fs statistics were significantly negative for DOM, MUS, and CAS, indicating an excess number of alleles, as expected in the case of recent population expansion. Additionally, Tajima’s D was significantly negative for all subspecies, including DOM, MUS, and CAS, further supporting the evidence of population size expansion. However, for BAC, Fu’s Fs and Tajima’s D were not statistically significant, indicating neutral selection and genetic drift. Mismatch distribution analysis for *M. musculus* (ALL) revealed multimodal curves, suggesting the presence of different genetic groups within the species. Harpending’s raggedness values were non-significant for all groups, indicating an expanding population. To examine the pattern of changes in effective population size, Bayesian skyline plot analyses were conducted. The effective population size of CAS and MUS gradually increased in the last 2,000 years, while the effective population size of DOM and BAC remained relatively constant until at least 1,000 years ago, and then decreased in the last century. Overall, *M. musculus* (ALL) displayed a constant effective population size between 11,000 and 2,500 years ago. From 2,000 years ago, there was a sharp increase in the effective population size (Fig. 5B). The beginning of population expansion in *M. musculus* (ALL) was estimated to have occurred approximately 155,000 years ago (Table 6).

**Fig. 5.**
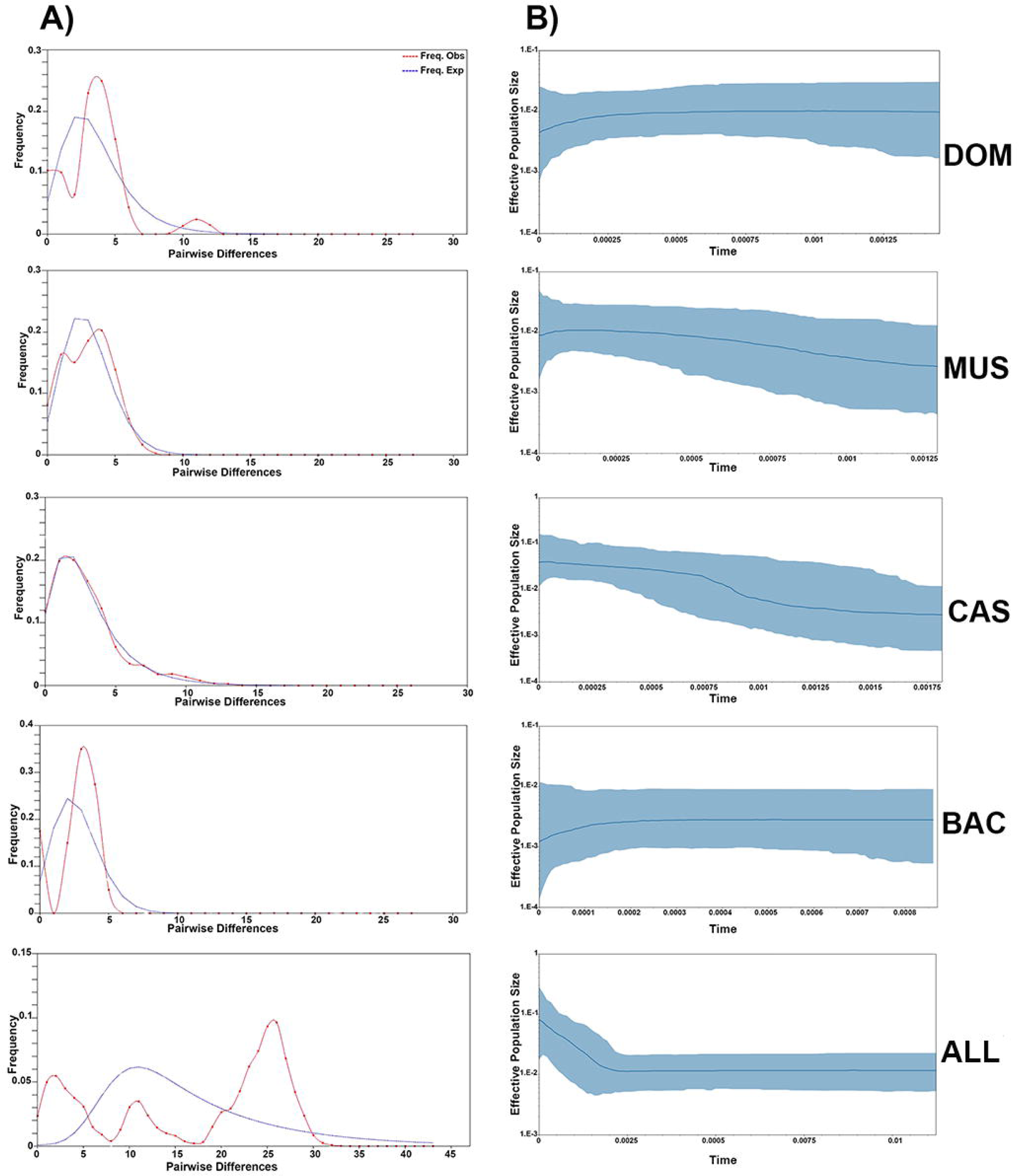
A) The mismatch distribution analysis (MMD); B) Bayesian skyline plot (BSP) analysis estimated based on the *Cytb* gene.

**Table 5.** Tajima’s D, Fu’s F and Harpending raggedness index for *Mus musculus* subspecies. Significant p values indicated with *.

|  | Tajima's D | Fu's Fs | Harpending raggedness<br>index |
| --- | --- | --- | --- |
| DOM | -1.82* | -5.39* | 0.05 |
| MUS | -1.60* | -19.55* | 0.01 |
| CAS | -1.91* | -24.83* | 0.02 |
| BAC | -0.20 | 1.24 | 0.13 |
| ALL | -0.84 | -23.65* | 0.00 |

**Table 6.** Tau (τ) values obtained from mismatch distribution analysis and the corresponding estimated time since expansion (t) for all groups. Three potential evolutionary rates were considered for these estimations. Notably, major estimated time since expansion values are highlighted in the table.

|  | Tau | t | t | t |
| --- | --- | --- | --- | --- |
|  |  | 0.11 | 0.047 | 0.028 |
| DOM | 3.94 | **19466** | 4555 | 7647 |
| MUS | 3.64 | **17984** | 4209 | 7065 |
| CAS | 1.36 | **6719** | 1572 | 2639 |
| BAC | 3.57 | 17638 | 4128 | 6929 |
| ALL | 7.99 | 39476 | 9239 | **15550** |

### Species Distribution Modelling

Models for Current Condition (CC) and Last Glacial Maximum (LGM) were run in 15 replicates (Fig. 6). The relative contribution of environmental variables for each period is presented in Table 7. The most influential environmental variables for the main models during the LGM were identified as follows: for DOM: Precipitation of Coldest Quarter (Bio_19), for MUS: Annual Mean Temperature (Bio_1), for CAS: Precipitation of Wettest Month (Bio_13), for BAC: Precipitation Seasonality (Bio_15), for ALL: Annual Mean Temperature (Bio_1). For the main models during the current conditions (CC): for DOM: Precipitation of Coldest Quarter (Bio_19), for MUS: Min Temperature of Coldest Month (Bio_6), for CAS: Precipitation of Wettest Month (Bio_13), for BAC: Mean Diurnal Range (Bio_2), for ALL: Min Temperature of Coldest Month (Bio_6). All models exhibited high predictive power, with very good AUC average values (Fig. 7). Habitat suitability for *M. musculus* exhibited changes from the last glacial maximum to the current period, with some regions losing suitable habitat. DOM transitioned from LGM to CC, losing suitable areas in East Asia but gaining new suitable areas in South Asia, particularly South India. MUS experienced a notable gain in new suitable areas from LGM to CC. CAS saw a minor loss of suitable areas in East Asia from LGM to CC. BAC, on the other hand, had more suitable habitats during the LGM than in the CC period.

**Fig. 6.**
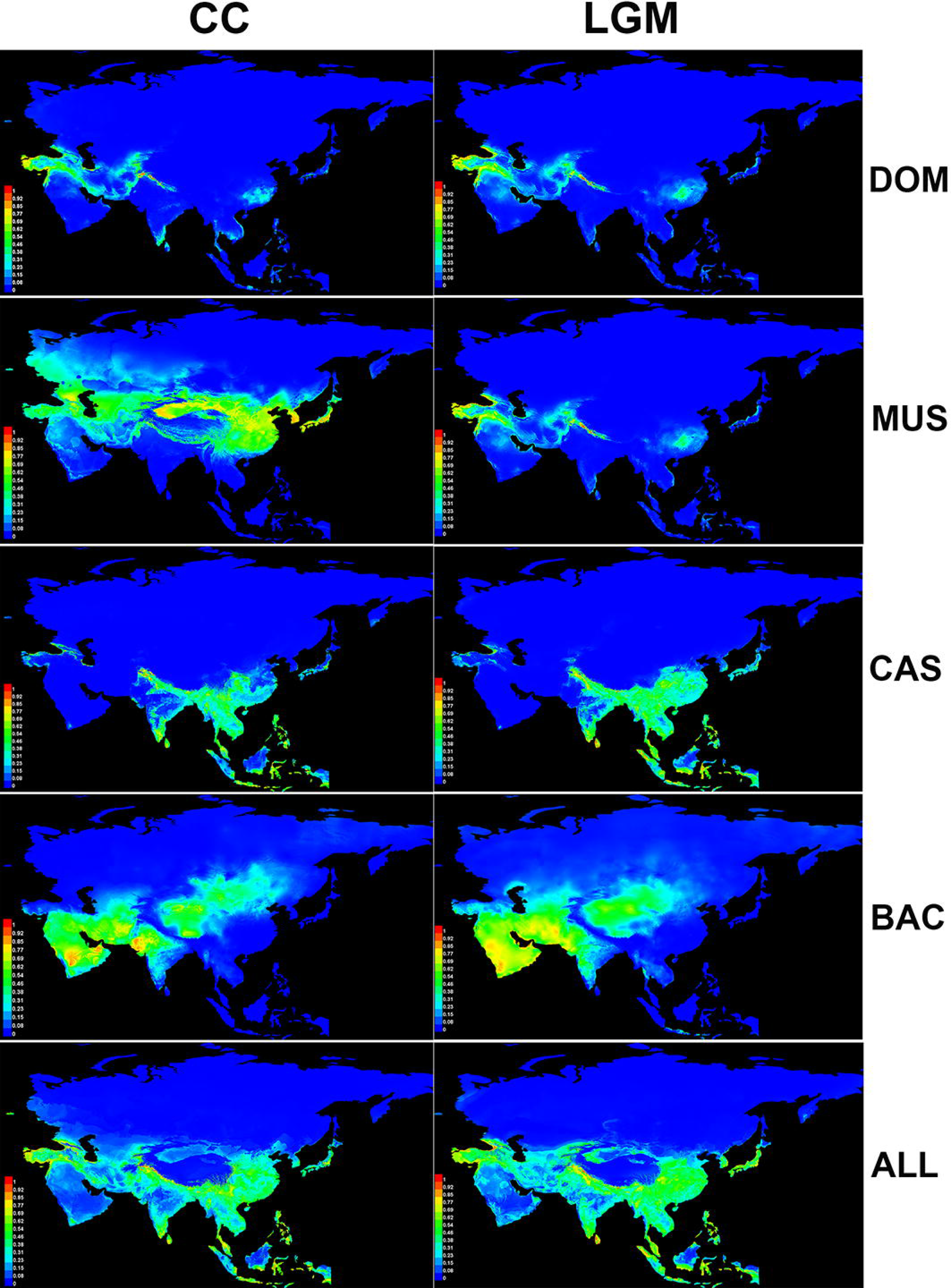
Species distribution modelling of *Mus musculus* for two distinct periods: Current Conditions (CC) and the Last Glacial Maximum (LGM). It visually represents changes in habitat suitability for the species over time.

**Fig. 7.**
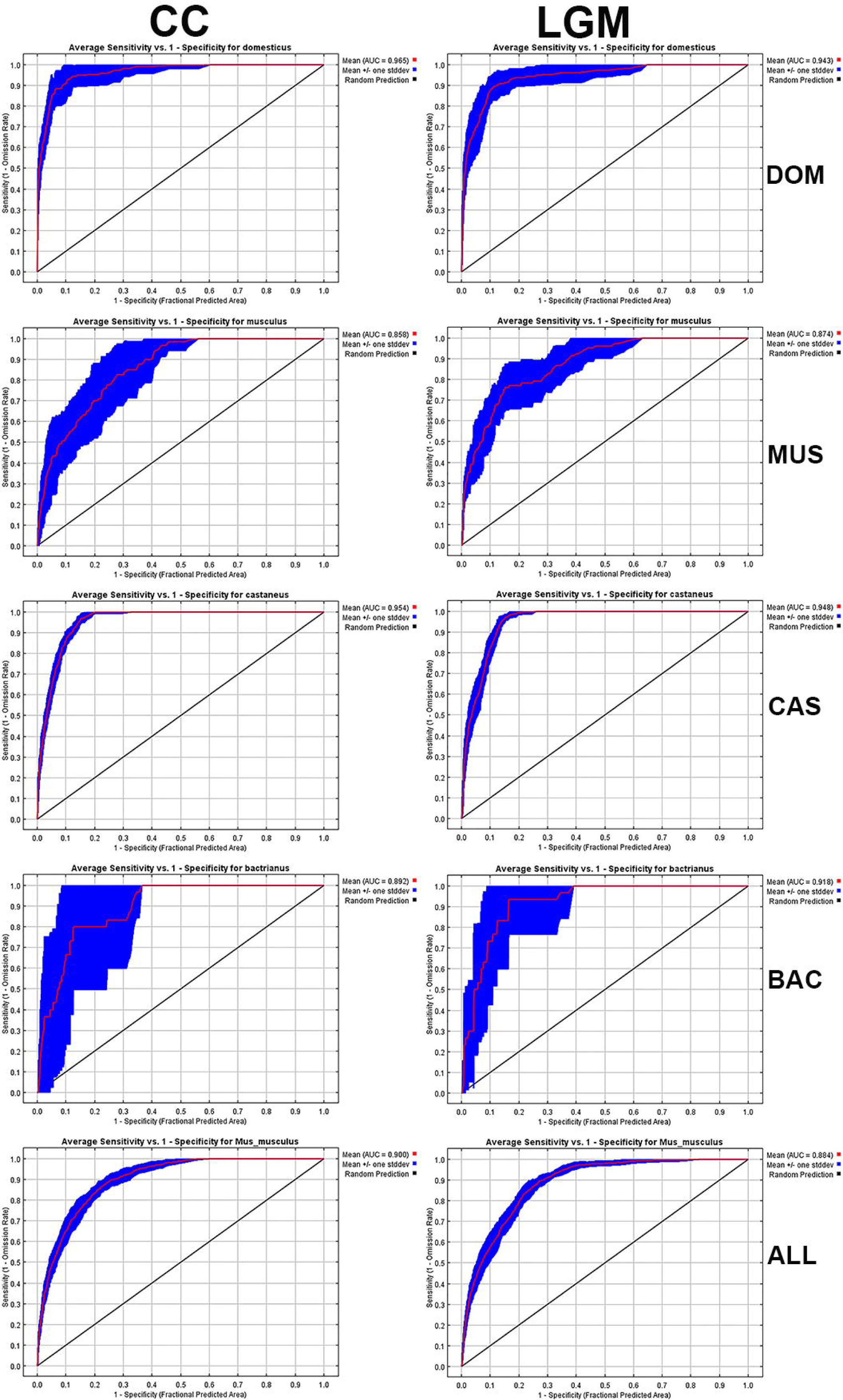
The Receiver Operating Characteristic (ROC) curve and AUC of *Mus musculus* in the Current Conditions (CC) and Last Glacial Maximum (LGM).

**Table 7.** Estimates of relative contributions of the environmental variables to the Maxent models, with the three major variables highlighted for each model.

| Variables | Current Condition |  |  |  |  | Last Glacial Maximum |  |  |  |  |
| --- | --- | --- | --- | --- | --- | --- | --- | --- | --- | --- |
|  | Percent Contribution |  |  |  |  | Percent Contribution |  |  |  |  |
|  | DOM | MUS | CAS | BAC | ALL | DOM | MUS | CAS | BAC | ALL |
| Bio_1 (Annual Mean Temperature) | 0.7 | **40.7** | 1 | 0 | **36.7** | 0 | **20.9** | 0.5 | 0 | **17.3** |
| Bio_2 (Mean Diurnal Range) | 0.4 | 1.5 | 0.5 | **18.9** | 1.2 | 1.2 | 2 | 1.6 | **40.8** | 1.5 |
| Bio_3 (Isothermality (BIO2/BIO7) ($\times 100$)) | **22** | 1.8 | 4.1 | 0 | 5.2 | 0.4 | 0 | 4.3 | 0 | 2.8 |
| Bio_4 (Temperature Seasonality) | 6.8 | 3 | **25.9** | 0.1 | 6.4 | 5 | 4.9 | 1.5 | 6 | 6.1 |
| Bio_5 (Max Temperature of Warmest Month) | 0 | 0.4 | 1.1 | 0 | 1.4 | 0 | 1.1 | 0.9 | 0 | 0.9 |
| Bio_6 (Min Temperature of Coldest Month) | 1.5 | 0.5 | **11.8** | 0 | 0.5 | **13.5** | **26.5** | **16.5** | 0 | **33.6** |
| Bio_7 (Temperature Annual Range (BIO5-BIO6)) | 0.5 | 1 | 0.2 | 0 | 0.3 | 0.7 | 4.3 | 1.5 | 0 | 2.5 |
| Bio_8 (Mean Temperature of Wettest Quarter) | **7.7** | 2 | 0.9 | 0.1 | 1.8 | **10.9** | 1.8 | 1 | 0 | 1.3 |
| Bio_9 (Mean Temperature of Driest Quarter) | 0.5 | **4.1** | 2.2 | **24.8** | 1 | 3.7 | 2.8 | 0.4 | **33.7** | 0.5 |
| Bio_10 (Mean Temperature of Warmest Quarter) | 0.4 | 0.7 | 0.6 | 0 | 1 | 0.1 | 0.4 | 1.5 | 0.2 | 2.8 |
| Bio_11 (Mean Temperature of Coldest Quarter) | 0.5 | **32.6** | 3.8 | 0 | **16.3** | 7.6 | **23.6** | 1.4 | 1.2 | 3.7 |
| Bio_12 (Annual Precipitation) | 0.4 | 0.3 | 2.2 | 11.4 | 1.5 | 0.4 | 0.1 | 0.4 | **8.3** | 1.1 |
| Bio_13 (Precipitation of Wettest Month) | 0.4 | 0.5 | **33.7** | 1.5 | 10.8 | 0.7 | 4.5 | **55.3** | 0.3 | **8.7** |
| Bio_14 (Precipitation of Driest Month) | 0.2 | 1.1 | 1.6 | 0 | 0.5 | 0.1 | 0.4 | 0.5 | 0 | 0.3 |
| Bio_15 (Precipitation Seasonality) | 1.1 | 3 | 0.5 | **40.2** | 1 | 0.5 | 0.1 | 1.3 | 2.8 | 1.7 |
| Bio_16 (Precipitation of Wettest Quarter) | 0 | 0 | 1.4 | 0.3 | 1.5 | 2 | 0.1 | 1.4 | 0.7 | 4 |
| Bio_17 (Precipitation of Driest Quarter) | 3.4 | 0.9 | 3.2 | 0.2 | 1.3 | 0.6 | 0.5 | 2.2 | 1.6 | 2.4 |
| Bio_18 (Precipitation of Warmest Quarter) | 2.9 | 2.5 | 3.8 | 1.9 | 0.7 | 0.7 | 4.6 | **5.4** | 4.3 | 0.9 |
| Bio_19 (Precipitation of Coldest Quarter) | **50.5** | 3.5 | 1.5 | 0.5 | **10.9** | **52** | 1.2 | 2.1 | 0.1 | 7.8 |

### Historical Biogeography

The historical biogeography analysis (Fig. 8) revealed that the divergence within the MRCA of all subspecies (node 101) likely originated in multiple biogeographic areas, including Broadleaf and Mixed Forests and Deserts and Xeric Shrublands. From these regions (West and Central Asia), a series of dispersal and vicariance events played a crucial role in the colonization of the species’ current ranges. Our analysis emphasized that dispersal events were more frequent within ecoregions, particularly in the Deserts and Xeric Shrublands. These intra-ecoregional dispersal events significantly contributed to the evolutionary history of the species, indicating that *M. musculus* primarily remained within its preferred habitat or similar ones. The divergence within the DOM lineage’s ancestor (node 54) was traced back to the Broadleaf and Mixed Forests. On the other hand, the ancestors of the BAC (node 55), CAS (node 81), and MUS (node 99) lineages had their origins in the Deserts and Xeric Shrublands ecoregion. Overall, the MRCA of all subspecies underwent a complex history, marked by 38 dispersal and 26 vicariance events, resulting in the expansion of their ranges and the emergence of distinct evolutionary lineages. Additionally, the MRCA of all subspecies experienced two extinction events, contributing to the intricate evolutionary landscape. The highest concentration of dispersal and vicariance events occurred approximately 0.11 Ma, coinciding with a period of increased divergence among subspecies. This temporal pattern indicated an accelerated rate of speciation and biogeographic differentiation. Our analysis revealed that most speciation events took place in the Deserts and Xeric Shrublands ecoregion, highlighting its significance as a hotspot for diversification. In contrast, the Montane Grasslands and Shrublands ecoregion played a comparatively minor role in the evolutionary history of *M. musculus* in Asia.

**Fig. 8.**
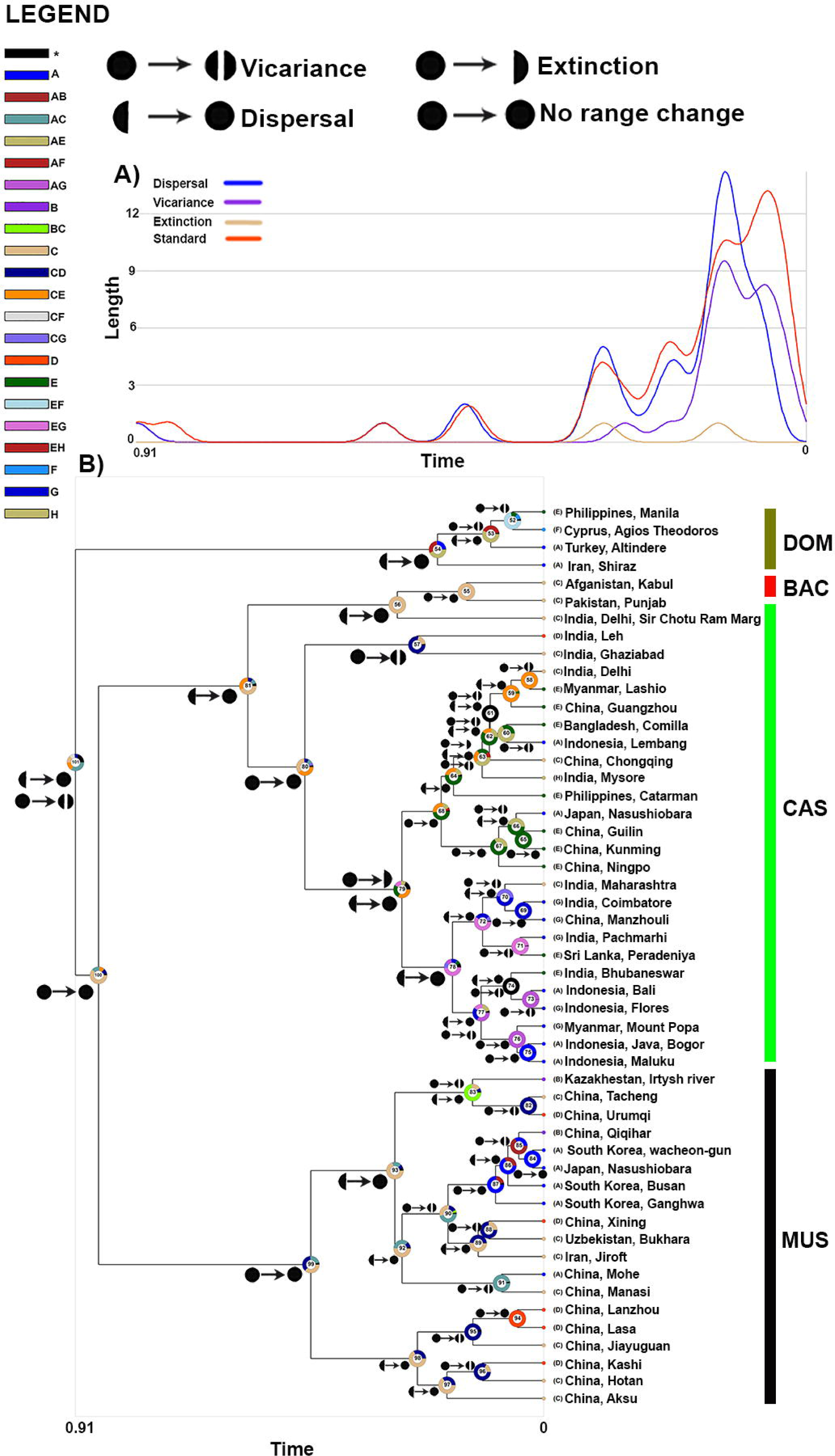
A) Diagram that shows the frequency of various ancestral origins (y-axis) against divergence time (x-axis); B) Historical biogeography analysis result, pie charts at internal nodes represent the marginal probabilities for each alternative ancestral area, and numbers represent node frequency percent.

## DISCUSSION

The house mouse, *Mus musculus*, has been a key model organism in evolutionary research, and its genetic diversity and phylogeography are of significant interest. In this study, we aimed to unravel the phylogeography and population genetic structure of *M. musculus* in Asian regions. Our analysis has shed light on various aspects of the species’ history, genetic diversity, and distribution patterns. Our analysis of mitochondrial markers, specifically the *Cytb* and *DL* regions, revealed a well-supported phylogenetic structure within the Asian *M. musculus* populations. These findings align with previous studies that have identified distinct subspecies within the species (Sakai et al., 2005; Frazer et al., 2007; Suzuki et al., 2013; Li et al., 2021; Amir Afzali & López-Antoñanzas, 2024). The monophyly of the four major subspecies; *Mus musculus musculus* (Eastern European house mouse), *Mus musculus domesticus* (Western European house mouse), *Mus musculus castaneus* (Southeastern Asian house mouse), and *Mus musculus bactrianus* (Southwestern Asian house mouse) suggests clear evolutionary lineages that have diverged over time. The timing of these divergences, as estimated from our analysis, provides insight into the evolutionary history of these subspecies. Our analyses identify *M. m. domesticus* as the sister group to *M. m. castaneus*, *M. m. bactrianus*, and *M. m. musculus*, with the highest genetic differentiation from the latter group. *M. m. castaneus* is the oldest subspecies, dating back approximately 0.3 Mya, while *M. m. bactrianus* is the youngest, originating around 0.1 Mya. These findings are well-supported in both Bayesian Inference (BI) and Maximum Likelihood (ML) analyses. Our historical biogeography analysis suggests that *M. m. bactrianus* originated in Northwest India, then expanded northward to North and Northeast Pakistan, Afghanistan, and East Iran. This pattern indicates peripheral speciation, with the Hindu Kush Mountains serving as a significant geographic barrier between *M. m. castaneus* and *M. m. bactrianus*. Species Distribution Modelling (SDM) and historical biogeography reveal distinct range expansion patterns. *M. m. musculus* and *M. m. domesticus* exhibit eastward expansion from the Last Glacial Maximum (LGM) to Current Conditions (CC), while *M. m. castaneus* and *M. m. bactrianus* experience westward expansion in Asia. These events may be linked to geological factors, including the penultimate glacial maximum (PGM) and subsequent rapid warming approximately 135,000 years ago, as well as human migration and agricultural practices (Li et al., 2021). Thus, the late Pleistocene global environmental fluctuations, occurring within the 100,000-year cycle, along with human activities, have played significant roles in the divergence events of *M. musculus*. Genetic diversity within these subspecies varies, with the highest diversity observed in *M. m. domesticus*, which may be attributed to its widespread distribution across multiple continents and the least diversity observed in *M. m. bactrianus* probably because it restricted in the Southwest Asia in Afghanistan and East Iran. The genetic divergence between subspecies, as indicated by pairwise genetic distances and *FST* values, underscores the significant differentiation among these groups, reflecting their historical isolation and limited gene flow. The Mantel test result supports the idea that geographical distance plays a crucial role in shaping genetic differentiation, highlighting the importance of geographical barriers in the diversification of *M. musculus* in Asia. The demographic history of *M. musculus* in Asia reveals interesting patterns. Fu’s Fs and Tajima’s D statistics suggest population expansions in the past, particularly in the *M. m. domesticus*, *M. m. musculus*, and *M. m. castaneus* subspecies, supported by multimodal mismatch distribution curves but genetic drift played an essential role in the shaping of *M. m. bactrianus* genetic structure. These findings align with the idea that the species has experienced demographic expansion during certain periods. Bayesian skyline plot analyses further illustrate population dynamics, showing variations in effective population sizes over time. The expansion of *M. m. castaneus* and *M. m. musculus* populations over the last 2,000 years, in contrast to the relatively stable or declining population sizes of *M. m. domesticus* and *M. m. bactrianus*, implies complex demographic histories for these subspecies. The estimation of the beginning of population expansion in *M. musculus* (ALL) at around 15,000 years ago coincides with the Pleistocene era, suggesting that climatic fluctuations during this period might have played a significant role in shaping the species’ demographic history. Further investigations into the specific drivers of these demographic patterns could provide valuable insights into the species’ response to historical environmental changes. Species distribution modelling provides valuable insights into the historical and current habitat suitability for *M. musculus* in Asia (Amir Afzali, 2024). The SDM results indicate changes in habitat suitability from the Last Glacial Maximum (LGM) to the Current Conditions (CC). These changes in habitat suitability can be attributed to climatic fluctuations and highlight the adaptability of the species to varying environmental conditions. For instance, *M. m. domesticus* expanded its range into South Asia, indicating its capacity to colonize new areas during favorable climatic periods. The importance of various environmental variables in driving these distribution patterns is evident, with temperature and precipitation variables playing key roles. Understanding these variables’ relative contributions can aid in predicting how *M. musculus* may respond to future environmental changes. These insights are valuable for conservation efforts and predicting potential range shifts in response to ongoing climate change. The historical biogeography analysis provides insights into the likely ancestral origins and the dispersal and vicariance events that have shaped the distribution of *M. musculus* in Asia. The West and Central Asian regions appear to be the likely points of origin for the MRCA of *M. musculus*, with subsequent dispersal events leading to colonization in various areas. The prevalence of dispersal within ecoregions and the high rate of divergence around 0.11 Ma suggest that geographical barriers and environmental factors have played a substantial role in driving speciation and distribution patterns. The predominance of speciation events in the Deserts and Xeric Shrublands ecoregion suggests that this habitat may have been a significant driver of differentiation within *M. musculus*. Further investigation into the ecological factors contributing to these patterns could yield a more comprehensive understanding of the species’ evolutionary history.

In conclusion, this study contributes to our understanding of the phylogeography and population genetic structure of *Mus musculus* in Asia. By using a combination of genetic data, historical demographic analysis, species distribution modelling, and historical biogeography, we have uncovered a complex history of diversification, demographic changes, and adaptation to changing environmental conditions. This knowledge not only enriches our understanding of this important model organism but also has broader implications for evolutionary and conservation biology. Future research could further investigate the specific environmental factors driving distribution patterns and demographic changes within each subspecies. Additionally, the integration of genomic data and the study of potential hybrid zones between subspecies could provide a more comprehensive view of the evolutionary dynamics of *M. musculus* in Asia.

## ACKNOWLEDGEMENT

I would like to express my sincere gratitude to Mohsen Amir Afzali for his invaluable assistance. Working alongside Mohsen has been a privilege, and I am truly thankful for his collaboration and friendship.

## DATA AVAILABILITY

All genetic sequences used in this study are publicly available in the GenBank and Supplementary Data files.

## SUPPLEMENTARY DATA

Supplementary data are available at *Journal of Mammalogy* online.

**Supplementary Data SD1.** Sampling localities of *M. musculus* in Asia.

**Supplementary Data SD2.** All *Cytb* sequences used for population genetic structure.

**Supplementary Data SD3.** All *DL* sequences.

**Supplementary Data SD4.** Concatenate data of *Cytb* and *DL* sequences.

**Supplementary Data SD5.** BI tree and posterior probabilities.

**Supplementary Data SD6.** ML tree and bootstrap values.

## LITERATURE CITED

Avise, J. C. (2000). Phylogeography: the history and formation of species. Harvard university press.

Amir Afzali, Y. (2024). Global climate change effect on Asian Mus musculus; Implication from last glacial maximum to the end of the 21st century. bioRxiv, 2024–03. 10.1101/2024.03.14.584992

Amir Afzali, Y., Darvish, J., & Yazdani-Moghaddam, F. (2017). Study of rodents’ fauna of the Jiroft, Kerman Province in southeast of Iran. Iranian Journal of Animal Biosystematics (IJAB), 13(1), 119–129. 10.22067/ijab.v13i1.59907

Amir Afzali, Y., & López-Antoñanzas, R. (2024). Molecular phylogeny and historical biogeography of Iranian murids (Rodentia: Muridae). Mammalian Biology, 104(1), 79–89. 10.1007/s42991-023-00390-3

Amir Afzali, Y., Naderloo, R., Keikhosravi, A., & Klaus, S. (2024). Geographic differentiation in the freshwater crab *Potamon persicum* Pretzmann, 1962 (Decapoda, Potamidae) in the Zagros Mountains: evidence from morphometry. Zoosystema, 46(4), 77–93. 10.5252/zoosystema2024v46a4

Amir Afzali, Y., Yazdani Moghaddam, F., Dianat, M., & Mahmoodi, A. (2018). Biosystematics Study of Golunda ellioti Gray, 1837 (Rodentia: Muridae) From Jiroft and Anbarabad Townships in Southeast of Iran. Journal of Research in Biology, 1. 1–5. 10.21859/jresbiol-e1522

Bouckaert, R., Vaughan, T. G., Barido-Sottani, J., Duchêne, S., Fourment, M., Gavryushkina, A., Heled, J., Jones, G., Kühnert, D., Maio, N. D., Matschiner, M., Mendes, F. K., Müller, N. F., Ogilvie, H. A., Plessis, L., Popinga, A., Rambaut, A., Rasmussen, D., Siveroni, I., Suchard, M. A., Wu, C. H., Xie, D., Zhang, C., Stadler, T., & Drummond, A. J., (2019). BEAST 2.5: An advanced software platform for Bayesian evolutionary analysis. PLoS computational biology, 15(4), e1006650. 10.1371/journal.pcbi.1006650

Church, D. M., Goodstadt, L., Hillier, L. W., Zody, M. C., Goldstein, S., She, X., Bult, C. J., Agarwala, R., Cherry, J. L., DiCuccio, M., Hlavina, W., Kapustin, Y., Meric, P., Maglott, D., Birtle, Z., Marques, A. C., Graves, T., Zhou, Sh., Teague, B., Potamousis, K., Churas, C., Place, M., Herschleb, J., Runnheim, R., Forrest, D., Amos-Landgraf, J., Schwartz, D. C., Cheng, Z., Lindblad-Toh, K., Eichler, E. E., Ponting, C. P., & Mouse Genome Sequencing Consortium. (2009). Lineage-specific biology revealed by a finished genome assembly of the mouse. PLoS biology, 7(5), e1000112. 10.1371/journal.pbio.1000112

Clement, M., Snell, Q., Walker, P., Posada, D., & Crandall, K. (2002). TCS: estimating gene genealogies. Parallel and Distributed Processing Symposium. In International Proceedings, 2(184): 10–1109.

Darriba, D., Taboada, G. L., Doallo, R., & Posada, D. (2012). jModelTest 2: more models, new heuristics and parallel computing. Nature methods, 9(8): 772–772. 10.1038/nmeth.2109

Darvish, J., Amirfazli, Y., & Hamidi, K. (2012). Further record of Golunda ellioti Gray, 1837 from South East of Iran with notes on its postcranial skeleton. Iranian Journal of Animal Biosystematics, 8(1). 10.22067/ijab.v8i1.25574

Edrisi, M., Rajabi-Maham, H., & Hashemian, N. (2018). Both Environment and Genetic Makeup Influence Sexual Behavior of House Mouse. Iranian Journal of Science and Technology, Transactions A: Science, 42(4), 1761–1769. 10.1007/s40995-018-0483-2

Elith, J., Graham, C. H., Anderson, R. P., Dudík, M., Ferrier, S., Guisan, A., Hijmans, R. J., Huettmann, F., Leathwick, J. R., Lehmann, A., Li, J., Lohmann, L. G., Loiselle, B. A., Manion, G., Moritz, C., Nakamura, M., Nakazawa, Y., Overton, J. M. M., Peterson, A. T., Phillips, S. J., Richardson, K., Scachetti-Pereira, R., Schapire, R. E., Soberón, J., Williams, S., Wisz, M. S., & Zimmermann, N. E. (2006). Novel methods improve prediction of species’ distributions from occurrence data. Ecography, 29: 129–151. 10.1111/j.2006.0906-7590.04596.x

Excoffier, L., & Lischer, H. E. (2010). Arlequin suite ver 3.5: a new series of programs to perform population genetics analyses under Linux and Windows. Molecular ecology resources, 10(3): 564–567. 10.1111/j.1755-0998.2010.02847.x

Frazer, K. A., Eskin, E., Kang, H. M., Bogue, M. A., Hinds, D. A., Beilharz, E. J., Gupta, R. V., Montgomery, J., Morenzoni, M. M., Nilsen, G. B., Pethiyagoda, C. L., Stuve, L. L., Johnson, F. M., Daly, M. J., C. M., & Cox, D. R. (2007). A sequence-based variation map of 8.27 million SNPs in inbred mouse strains. Nature, 448(7157), 1050–1053. 10.1038/nature06067

Guénet, J. L., & Bonhomme, F. (2003). Wild mice: an ever-increasing contribution to a popular mammalian model. Trends in Genetics, 19(1), 24–31.

Hall, T. A. (1999). BioEdit: a user-friendly biological sequence alignment editor and analysis program for Windows 95/98/NT. In Nucleic acids symposium series (Vol. 41, No. 41, pp. 95–98).

Hardouin, E. A., Orth, A., Teschke, M., Darvish, J., Tautz, D., & Bonhomme, F. (2015). Eurasian house mouse (*Mus musculus* L.) differentiation at microsatellite loci identifies the Iranian plateau as a phylogeographic hotspot. BMC evolutionary biology, 15(1), 1–12. 10.1186/s12862-015-0306-4

Hasegawa, M., Kishino, H., Yano, T. (1985). Dating of the Human-Ape Splitting by a molecular Clock of Mitochondrial DNA. Journal of Molecular Evolution, 22, 160–174. 174. 10.1007/BF02101694

Hashemian, N., Rajabi-Maham, H., & Edrisi, M. (2017). Genetic vs environment influences on house mouse hybrid zone in Iran. Journal of Genetic Engineering and Biotechnology, 15:483–488. 10.1016/j.jgeb.2017.06.002

Hernandez, P. A., Graham, C. H., Master, L. L., & Albert, D. L. (2006). The effect of sample size and species characteristics on performance of different species distribution modeling methods. Ecography, 29: 773–785. 10.1111/j.0906-7590.2006.04700.x

Hijmans, R. J., Cameron, S. E., Parra, J. L., Jones, P. G., & Jarvis, A. (2005). Very high-resolution interpolated climate surfaces for global land areas. International Journal of Climatology: A Journal of the Royal Meteorological Society, 25(15): 1965–1978. 10.1002/joc.1276

Honda, A., Murakami, S., Harada, M., Tsuchiya, K., Kinoshita, G., & Suzuki, H. (2019). Late Pleistocene climate change and population dynamics of Japanese Myodes voles inferred from mitochondrial cytochrome b sequences. Journal of Mammalogy, 100(4), 1156–1168. 10.1093/jmammal/gyz093

Kumar, S., Stecher, G., Li, M., Knyaz, C., & Tamura, K. (2018). MEGA X: molecular evolutionary genetics analysis across computing platforms. Molecular biology and evolution, 35(6), 1547. 10.1093%2Fmolbev%2Fmsy096

Li, Y., Fujiwara, K., Osada, N., Kawai, Y., Takada, T., Kryukov, A. P., Abe, K., Yonekawa, H., Shiroishi, T., Moriwaki, K., Saitou, N., & Suzuki, H. (2021). House mouse Mus musculus dispersal in East Eurasia inferred from 98 newly determined complete mitochondrial genome sequences. Heredity, 126(1), 132–147. 10.1038/s41437-020-00364-y

Maltsev, A. N., Stakheev, V. V., & Kotenkova, E. V. (2016). Role of invasions in formation of phylogeographic structure of house mouse (*Mus musculus*) in some areas of Russia and the near abroad. Russian journal of biological invasions, 7, 255–267. 10.1134/S2075111716030061

Matzke N (2013) BioGeoBEARS: BioGeography with Bayesian (and Likelihood) Evolutionary Analysis in R Scripts. University of California, Berkeley, Berkeley, CA

Maung Maung Theint, S., Thwe, T., Myat Myat Zaw, K., Shimada, T., Bawm, S., Kobayashi, M., Saing, K. M., Katakura, K., Arai, S., & Suzuki, H. (2021). Late quaternary environmental and human impacts on the mitochondrial DNA diversity of four commensal rodents in Myanmar. Journal of Mammalian Evolution, 28, 497–509. 10.1007/s10914-020-09519-4

Mitchell-Jones, A. J., Amori, G., Bogdanowicz, W., Krystufek, B., Reijnders, P. J. H., Spitzenberger, F., Stubbe, M., Thissen, J. B. M., & Zima, J. E. (1999). The atlas of European mammals (Vol. 3). London: Academic Press.

Montgelard, C., Bentz, S., Tirard, C., Verneau, O., & Catzeflis, F. M. (2002). Molecular systematics of Sciurognathi (Rodentia): the mitochondrial cytochrome *b* and 12S rRNA genes support the Anomaluroidea (Pedetidae and Anomaluridae). Molecular Phylogenetics and Evolution, 22(2), 220–233. 10.1006/mpev.2001.1056

Musser, G. G., & Carleton, M. D. (2005). “Superfamily Muroidea”. In Wilson, D. E., & Reeder, D. M. (eds.). Mammal Species of the World: A Taxonomic and Geographic Reference (3rd ed.). Baltimore: Johns Hopkins University Press. pp. 894–1531. ISBN 978-0-8018–8221-0.

Olson, D. M., Dinerstein, E., Wikramanayake, E. D., Burgess, N. D., Powell, G. V., Underwood, E. C., D’amico, J. A., Itoua, I., Strand, E. E., Morrison, J. C., Loucks, C. J., Allnutt, T. F., Ricketts, T. H., Kura, Y., Lamoreux, J. F., Wettengel, W. W., Hedao, P., & Kassem, K. R. (2001). Terrestrial Ecoregions of the World: A New Map of Life on Earth: A new global map of terrestrial ecoregions provides an innovative tool for conserving biodiversity. BioScience, 51:933–938. 10.1641/0006-3568(2001)051[0933:TEOTWA]2.0.CO;2

Phillips, S. J., Anderson, R. P., & Schapire, R. E. (2006). Maximum entropy modeling of species geographic distributions. Ecological modelling, 190(3–4): 231–259. 10.1016/j.ecolmodel.2005.03.026

Rambaut, A. (2018). FigTree v.1.4.4. Available at: http://tree.bio.ed.ac.uk/software/figtree/.

Ree, R. H., Moore, B. R., Webb, C. O., & Donoghue, M. J. (2005). A likelihood framework for inferring the evolution of geographic range on phylogenetic trees. Evolution, 59(11), 2299–2311. 10.1111/j.0014-3820.2005.tb00940.x

Ree, R. H., & Sanmartín, I. (2018). Conceptual and statistical problems with the DEC+ J model of founderDevent speciation and its comparison with DEC via model selection. Journal of Biogeography, 45(4), 741–749. 10.1111/jbi.13173

Rhoden, C. M., Peterman, W. E., & Taylor, C. A. (2017). Maxent-directed field surveys identify new populations of narrowly endemic habitat specialists. PeerJ, 5, e3632. 10.7717/peerj.3632

Rogers, A. R. (1995). Genetic evidence for a Pleistocene population explosion. Evolution, 49(4), 608–615. 10.1111/j.1558-5646.1995.tb02297.x

Rogers, A. R., & Harpending, H. (1992). Population growth makes waves in the distribution of pairwise genetic differences. Molecular biology and evolution, 9(3), 552–569. 10.1093/oxfordjournals.molbev.a040727

Rozas, J., Ferrer-Mata, A., Sánchez-DelBarrio, J. C., Guirao-Rico, S., Librado, P., Ramos-Onsins, S. E., & Sánchez-Gracia, A. (2017). DnaSP 6: DNA sequence polymorphism analysis of large data sets. Molecular biology and evolution, 34(12), 3299–3302. 10.1093/molbev/msx248

Sakai, T., Kikkawa, Y., Miura, I., Inoue, T., Moriwaki, K., Shiroishi, T., Satta, Y., Takahata, N., & Yonekawa, H. (2005). Origins of mouse inbred strains deduced from whole-genome scanning by polymorphic microsatellite loci. Mammalian Genome, 16, 11–19. 10.1007/s00344-004-3013-9

Shimada, T., Aplin, K. P., & Suzuki, H. (2010). *Mus lepidoides* (Muridae, Rodentia) of central Burma is a distinct species of potentially great evolutionary and biogeographic significance. Zoological science, 27(5), 449–459. 10.2108/zsj.27.449

Sillero, N., & Carretero, M. A. (2012). Modelling the past and future distribution of contracting species. The Iberian lizard *Podarcis carbonelli* (Squamata: Lacertidae) as a case study. Zoologischer Anzeiger, 252, 289–298. 10.1016/j.jcz.2012.08.004

Stamatakis, A. (2014). RAxML version 8: a tool for phylogenetic analysis and post-analysis of large phylogenies. Bioinformatics, 30(9), 1312–1313. 10.1093%2Fbioinformatics%2Fbtu033

Suzuki, H., Nunome, M., Kinoshita, G., Aplin, K. P., Vogel, P., Kryukov, A. P., Jin, M. L., Han, S. H., Maryanto, I., Tsuchiya, K., Ikeda, H., Shiroishi, T., Yonekawa, H., & Moriwaki, K. (2013). Evolutionary and dispersal history of Eurasian house mice *Mus musculus* clarified by more extensive geographic sampling of mitochondrial DNA. Heredity, 111(5), 375–390. 10.1038/hdy.2013.60

Suzuki, H., Shimada, T., Terashima, M., Tsuchiya, K., & Aplin, K. (2004). Temporal, spatial, and ecological modes of evolution of Eurasian *Mus* based on mitochondrial and nuclear gene sequences. Molecular phylogenetics and evolution, 33(3), 626–646. 10.1016/j.ympev.2004.08.003

Tavaré, S. (1986). Some probabilistic and statistical problems on the analysis of DNA sequence. Lecture of Mathematics for Life Science, 17, 57.

Yasuda, S. P., Vogel, P., Tsuchiya, K., Han, S. H., Lin, L. K., & Suzuki, H. (2005). Phylogeographic patterning of mtDNA in the widely distributed harvest mouse (*Micromys minutus*) suggests dramatic cycles of range contraction and expansion during the mid-to late Pleistocene. Canadian Journal of Zoology, 83(11), 1411–1420. 10.1139/z05-139

